# Evolutionary dynamics of chloroplast genomes in low light: a case study of the endolithic green alga Ostreobium quekettii

**DOI:** 10.1101/049833

**Authors:** Vanessa R. Marcelino, Ma Chiela M. Cremen, Chistopher J. Jackson, Anthony W.D. Larkum, Heroen Verbruggen

**Author notes:** **Author for correspondence:** Vanessa Rossetto, Marcelino School of Biosciences (bldg. 122, room 101), University of Melbourne, VIC 3010, Australia.

## Abstract

Some photosynthetic organisms live in extremely low light environments. Light limitation is associated with selective forces as well as reduced exposure to mutagens, and over evolutionary timescales it can leave a footprint on species’ genome. Here we present the chloroplast genomes of four green algae (Bryopsidales, Ulvophyceae), including the endolithic (limestone-boring) alga *Ostreobium quekettii*, which is a low light specialist. We use phylogenetic models and comparative genomic tools to investigate whether the chloroplast genome of *Ostreobium* corresponds to our expectations of how low light would affect genome evolution. *Ostreobium* has the smallest and most gene-dense chloroplast genome among Ulvophyceae reported to date, matching our expectation that light limitation would impose resource constraints. Rates of molecular evolution are significantly slower along the phylogenetic branch leading to *Ostreobium*, in agreement with the expected effects of low light and energy levels on molecular evolution. Given the exceptional ability of our model organism to photosynthesize under extreme low light conditions, we expected to observe positive selection in genes related to the photosynthetic machinery. However, we observed stronger purifying selection in these genes, which might either reflect a lack of power to detect episodic positive selection followed by purifying selection and/or a strengthening of purifying selection due to the loss of a gene related to light sensitivity. Besides shedding light on the genome dynamics associated with a low light lifestyle, this study helps to resolve the role of environmental factors in shaping the diversity of genome architectures observed in nature.

**Data deposition:** Chloroplast genome sequences will be deposited in GenBank

## Introduction

Light is rapidly attenuated underwater, yet some photosynthetic organisms thrive in extremely low light marine habitats (Shashar and Stambler 1992; Hawes and Schwarz 1999; Mock and Kroon 2002; Larkum et al. 2003). Specialized lifestyles may leave a footprint on organisms’ genomes (Dutta and Paul 2012; Raven et al. 2013). For example, high-light and low light strains of the cyanobacterium *Prochlorococcus* have different genome sizes, GC contents and rates of molecular evolution, among other genome features that have been associated with their niche specialization (Hess et al. 2001; Rocap et al. 2003; Dufresne et al. 2005; Paul et al. 2010). Similar genome studies targeting eukaryotic algae have only begun to emerge in recent times (see Raven et al. 2013 for a review). Different ecotypes of the microalga *Ostreococcus*, for example, show distinctive genome traits (Jancek et al. 2008), although in this case it is not clear whether low light has played a role.

Light modulates the production of high-energy cofactors (ATP and NADPH) and the uptake of nitrogen (MacIsaac and Dugdale 1972; Cochlan et al. 1991; Kirk 1994), therefore it is logical to expect that the genome architecture of lineages living under low light conditions is influenced by resource constraints. It has been proposed that the loss of unessential genes, intergenic spacers and introns (genome streamlining) may be driven by selection for diminishing resources and time required for replication and growth (Giovannoni et al. 2005; Hessen et al. 2010; Wolf and Koonin 2013). Genome architecture can also be affected by limited supply of key elements such as nitrogen and phosphorus: different nucleotides and amino acids differ in their atomic composition, so molecules containing less atoms of the limiting nutrient may provide a selective advantage in certain niches (Acquisti et al. 2009; Elser et al. 2011; Raven et al. 2013). The *Prochlorococcus* strain with the smallest genome and highest content of nitrogen-poor molecules is found in surface waters, where irradiance is higher but nutrients are more depleted than in the habitat of the low light strain (Rocap et al. 2003; Dufresne et al. 2005). One could expect that when light is low enough to restrict growth rates and nitrogen uptake, organisms with small genomes and a high proportion of nitrogen-poor molecules may have better evolutionary fitness.

Sunlight may leave footprints in a genome by directly or indirectly altering molecular rates of evolution (the molecular pacemaker). Light is a major contributor to environmental energy including solar radiation, thermal energy and chemical (metabolic) energy (Clarke and Gaston 2006). Environmental energy stimulates metabolism at many levels, and it is known that energy-rich habitats lead to higher evolutionary rates (Davies et al. 2004; Clarke and Gaston 2006). Solar radiation, especially ultraviolet (UV), also plays a direct mutagenic role and may thus accelerate molecular evolution (Rothschild 1999; Willis et al. 2009). Thermal and chemical energy also depend on light: light incidence increases temperatures (e.g. in the tropics) and supports primary productivity (and consequently increases the energy available for metabolism and growth). Oxidative DNA damage generally occurs during metabolic reactions, therefore higher metabolic rates can lead to higher mutation rates (Gillooly et al. 2005). Generation times also play into it, being shorter and fixing mutations (on populations) more rapidly when the environmental energy is higher, which often happens when there is a combination of higher temperatures, metabolic rates and solar radiation (Rohde 1992; Wright and Rohde 2013). Therefore it is reasonable to expect that organisms living in low-energy areas, like shaded habitats, have slower rates of molecular evolution.

Challenging environments may impose particular selective regimes, which could leave a footprint of positive selection in genes undergoing adaptation (e.g. Qu et al. 2013). For example, evidence of positive selection in the Rubisco gene (involved in carbon fixation) in mosses has been associated with its adaptation to the declining levels of atmospheric CO_2_ since their origination in the Ordovician (Raven and Colmer 2016). In cases of organisms living in extremely low light, it would be reasonable to expect positive selection in genes related to the photosynthetic machinery. To our knowledge, this has never been tested in eukaryotic algae.

The siphonous green alga *Ostreobium* is a convenient organism to investigate photosynthesis under low light conditions (Fork and Larkum 1989; Koehne et al. 1999; Wilhelm and Jakob 2006). *Ostreobium* has an endolithic (limestone-boring) lifestyle: it bores into carbonate substrates and populates all sorts of marine limestones worldwide, including shells and coral skeletons. Only a small proportion of the available light reaches *Ostreobium* in its usual habitat: about 99% of the light is attenuated by the first millimeter of limestone (Nienow et al. 1988; Matthes et al. 2001). Other photosynthetic organisms living on the limestone substrate can further attenuate light: the living tissue of corals and their zooxanthellae, for example, absorb 95 – 99.9% of the available light (Halldal 1968; Schlichter et al. 1997). Even under these extreme low light conditions, *Ostreobium* carries out oxygenic photosynthesis (Kühl et al. 2008). It has been shown that some low light cyanobacteria enhance their light interception by manufacturing special near-infrared (NIR) absorbing chlorophylls (Chl *d* and *ƒ*; Chen and Blankenship 2011). Therefore it is of interest that *Ostreobium* species coexist in a similar habitat, where they must have special mechanisms for surviving under such low light NIR-enriched conditions (Magnusson et al. 2007). Another low light strategy in *Ostreobium* also allows it to grow in deep waters, where it is abundant even at depths over 200 meters, and where only a handful of algal species can persist (Littler et al. 1985; Dullo et al. 1995; Aponte and Ballantine 2001). Here the light is filtered strongly in oceanic water with a peak at around 470-480 nm (Larkum and Barrett 1983) and a different light harvesting strategy is employed: the carotenoid siphonaxanthin transfers light energy to chlorophyll and the reaction centers (Kageyama et al. 1977). Thus the success of *Ostreobium* in terms of its cosmopolitan distribution is associated not only with its efficiency in light utilization and but also its ability to employ a range of light harvesting strategies (Fork and Larkum 1989; Schlichter et al. 1997; Tribollet 2008), for which the underlying genomic basis has never been explored.

The light-driven genomic traits of *Ostreobium* can only be investigated in a comparative framework. While algal nuclear genome sequences are still scarce, chloroplast genomes are better sampled and constitute a powerful tool for molecular evolutionary studies (Wicke and Schneeweiss 2015). *Ostreobium* belongs to the Bryopsidales (Ulvophyceae), a diverse order of seaweeds for which only four chloroplast genomes are available. Additional chloroplast genomes of species living in higher-light habitats, especially those from early-diverging lineages within the order, will help us investigate such genomic traits.

The goal of this study is to evaluate the evolutionary dynamics of the chloroplast genome of the low light alga *Ostreobium* using comparative and phylogenetic methods. Because comparative analyses in a phylogenetic context require a sufficiently large sample of genomes, we present the chloroplast genomes of four green algae, including *Ostreobium quekettii* and members from three other families in the same order, all previously unsequenced. We use a combination of stoichiogenomics (the study of elemental composition of macromolecules (Elser et al. 2011)) and models of molecular rate variation to investigate our expectations for a lineage adapted to low light conditions. Our first expectation relates to light-dependent resource limitation: if the *Ostreobium* lineage has evolved in low-energy and low-nutrient conditions, its chloroplast genome can be expected to be smaller, more compact (i.e. with less intergenic spacers, introns and repeats) and contain less nitrogen than the chloroplast genomes of related algae. Our second expectation is that the phylogenetic branch leading to *Ostreobium* has slower rates of molecular evolution than other branches in the phylogeny as a consequence of the presumably fewer mutations induced by UV and/or metabolism, and slower generation times imposed by its low energy niche. Lastly, based on *Ostreobium*’s highly efficient light utilization, we would expect genes related to its photosynthetic machinery to have experienced positive selection.

## Materials and Methods

### Samples and Sequencing

The *Ostreobium quekettii* sample was obtained from the Culture Collection of Algae at the University of Gottingen (SAG 6.99). The *Halimeda discoidea* sample (voucher HV04923) was collected in New Ireland, Papua New Guinea in 2014. *Caulerpa cliftonii* (voucher HV03798) was collected in Point Lonsdale (VIC, Australia) in 2013. The strain of *Derbesia sp*. (WEST4838) was provided by John West (University of Melbourne). Total genomic DNA was extracted using a modified cetyl trimethylammonium bromide (CTAB) method as described in Cremen et al. (2016). *Ostreobium* was sequenced in an Illumina NextSeq 500 platform at the Georgia Genome Facility, while the other samples were sequenced on the Illumina HiSeq platform at the Genome Center of the Cold Spring Harbor Marine Laboratory. Sequences were submitted to GenBank (accession numbers to be provided).

### Assembly and Annotation

Sequences were assembled using CLC Genomics Workbench 7.5.1 (http://www.clcbio.com). Circularity were resolved by comparing the CLC assembly with assemblies generated independently with MEGAHIT (Li et al. 2015) and SOAPdenovo2 (Luo et al. 2012). Scaffolds were resolved with GapCloser (Luo et al. 2012) and by mapping the sequence reads to the scaffold in Geneious 9.0.4 (Kearse et al. 2012). Coverage was also calculated with Geneious.

We used a combination of automated pipelines and manual editing to annotate the plastomes. Sequences were submitted to the MFannot (http://megasun.bch.umontreal.ca/RNAweasel/), DOGMA (Wyman et al. 2004) and ARAGORN (Laslett 2004) online tools. The annotations were manually compared in Geneious and added to the final annotation track once their accuracy was verified. Intron types were determined with RNAweasel (http://megasun.bch.umontreal.ca/RNAweasel/). Start and stop positions of coding sequences and introns were visually verified by aligning them with sequences of other algae using MAFFT (Katoh et al. 2002).

In order to compare the genome of *Ostreobium* with other Ulvophyceae, the plastomes of *Bryopsis plumosa* (NC_026795), *Tydemania expeditions* (NC_026796), *Ulva* sp. (KP720616), *Pseudendoclonium akinetum* (AY835431) and *Oltmannsiellopsis viridis* (NC_008099) were downloaded from GenBank. The genome features from all analyzed plastomes were obtained using Geneious. We excluded all hypothetical ORFs with less than 300bp and re-annotated the *tilS* gene as a pseudogene (not a CDS) in species where it has a frame shift or a stop codon in the middle of the gene. The Geneious implementation of Phobos v.3.3.11 (Mayer 2007) was used to identify tandem repeats with lengths between 15 and 1000bp, using the “perfect” search mode. Nitrogen (N) content was quantified based on the counts of N atoms per nucleotide or amino acid using the formula described in Acquisti et al. (2009): Σ.(*n*_*i*_×*p*_*i*_) where *n*_*i*_ is the number of N atoms in the *i*-th base and *p*_*i*_ is the proportion of each base in the plastome. For the nucleotide counts, *n*_C_ = *n*_G_ = 4 and *n*_A_ = *n*_T_ = 3.5 (Acquisti et al. 2009). For the transcribed genomic segments (transcriptome) we used *n*_A_ = 5, *n*_T_ = 2, *n*_G_ = 5, and *n*_C_ = 3. For the amino acids counts we used *n* = 2 for asparagine, glutamine, lysine and tryptophan; *n* = 3 for histidine; *n* = 4 for arginine; and *n* = 1 for other amino acids.

### Phylogeny, rates of evolution and selection analysis

We aligned the coding sequences of all species at the amino acid level using MAFFT (Katoh et al. 2002) and then converted back to nucleotides using RevTrans (Wernersson and Pedersen 2003). In addition to *tilS, the ftsH, rpoB, rpoC1 rpoC2* and *ycf1* genes could not be reliably aligned and were excluded from downstream analyses. A Maximum Likelihood phylogeny was built using RAxML (Stamatakis 2006) with the GTR+Γ model, a partitioning strategy separating 1^st^, 2^nd^ and 3^rd^ codon positions, and a rapid bootstrap search of 500 replicates. *Oltmannsiellopsis viridis, Pseudendoclonium akinetum* and *Ulva* sp. were used as outgroups.

We studied lineage-specific rates of evolution using the baseml program from the PAML v.4.7 package (Yang 2007). The fit of a model with unique rates of evolution across all branches (global clock) was compared to a model with a different rate for the *Ostreobium* lineage (local clock). The Akaike Information Criterion (AIC) was used to compare model fit.

To evaluate whether photosynthetic genes have been under positive selection in the *Ostreobium* lineage, we grouped (concatenated) genes into 15 gene classes as defined in Wicke et al. (2011). Individual gene alignments containing less than 4 species were excluded from the analyses. We tested the fit of a model with differential *d*_N_/*d*_S_ ratio (ω) for *Ostreobium* and the background lineages, versus a model with a universal ω for all branches (the null hypothesis), using the codeml program implemented in PAML (Yang 1998; Yang 2007). We used the F3×4 codon model and model fit was compared with AIC for each gene (Table 2). We also used the random effects branch-site model implemented in HyPhy (Kosakovsky Pond et al. 2005; Kosakovsky Pond et al. 2011). This method allows detecting the timing of positive selection in a phylogeny and the proportion of sites under selection. Positive selection in the *Ostreobium* lineage was evaluated by a Likelihood ratio test.

## Results

### Four new chloroplast genomes of Bryopsidales

The chloroplast genomes of *Ostreobium queketti*, *Halimeda discoidea, Derbesia* sp. and *Caulerpa cliftonii* were sequenced. The *Ostreobium* chloroplast genome was recovered as a complete (circular mapping) molecule with a mean coverage of 235×. The mean coverage was high for *Halimeda* (3,983×), but two scaffolds had a very low coverage and could not be resolved with confidence. For *Caulerpa*, the mean coverage was 1,116×, but this species also had one scaffold that could not be determined with certainty. These unresolved scaffolds in *Caulerpa* and *Halimeda* likely represent a polymorphic number of repeats. The overall high coverage and the presence of the great majority of plastid genes (Supplementary table 1) indicate that we likely recovered sequences for all genes in the chloroplast and the few non-coding regions that have not been resolved should not interfere in our analysis. We also obtained the complete chloroplast genome of *Derbesia* sp., with a mean coverage of 469×. Genome sizes and main features are reported in Table 1.

Gene content of *Ostreobium* is similar to related algae but it lacks the *chloroplast envelope membrane protein* gene (*cemA*) that is present in all other Ulvophyceae sequenced to date (Supplementary table 1). The *tRNA(Ile)-lysidine synthase* gene (*tilS*) seems to be a pseudogene in *Ostreobium*, *Halimeda* and *Derbesia* as it contains multiple in-frame stop codons. We could not identify it at all in *Caulerpa cliftonii* (i.e. no tblastx hits with e-values < 0.001 and identity > 50%, using *Bryopsis plumosa* genes as reference), although this pseudogene has been found in another *Caulerpa* species (Zuccarello et al. 2009). None of our chloroplast genomes have the *organelle division inhibitor factor* gene (*minD*), supporting the suggestion that this gene has been lost from the chloroplasts of Bryopsidales (Leliaert and Lopez-Bautista 2015). Despite an overall highly conserved gene content, the Ulvophyceae genomes have multiple rearrangements as indicated in the Mauve alignment (Figure 1).

**Figure 1:**
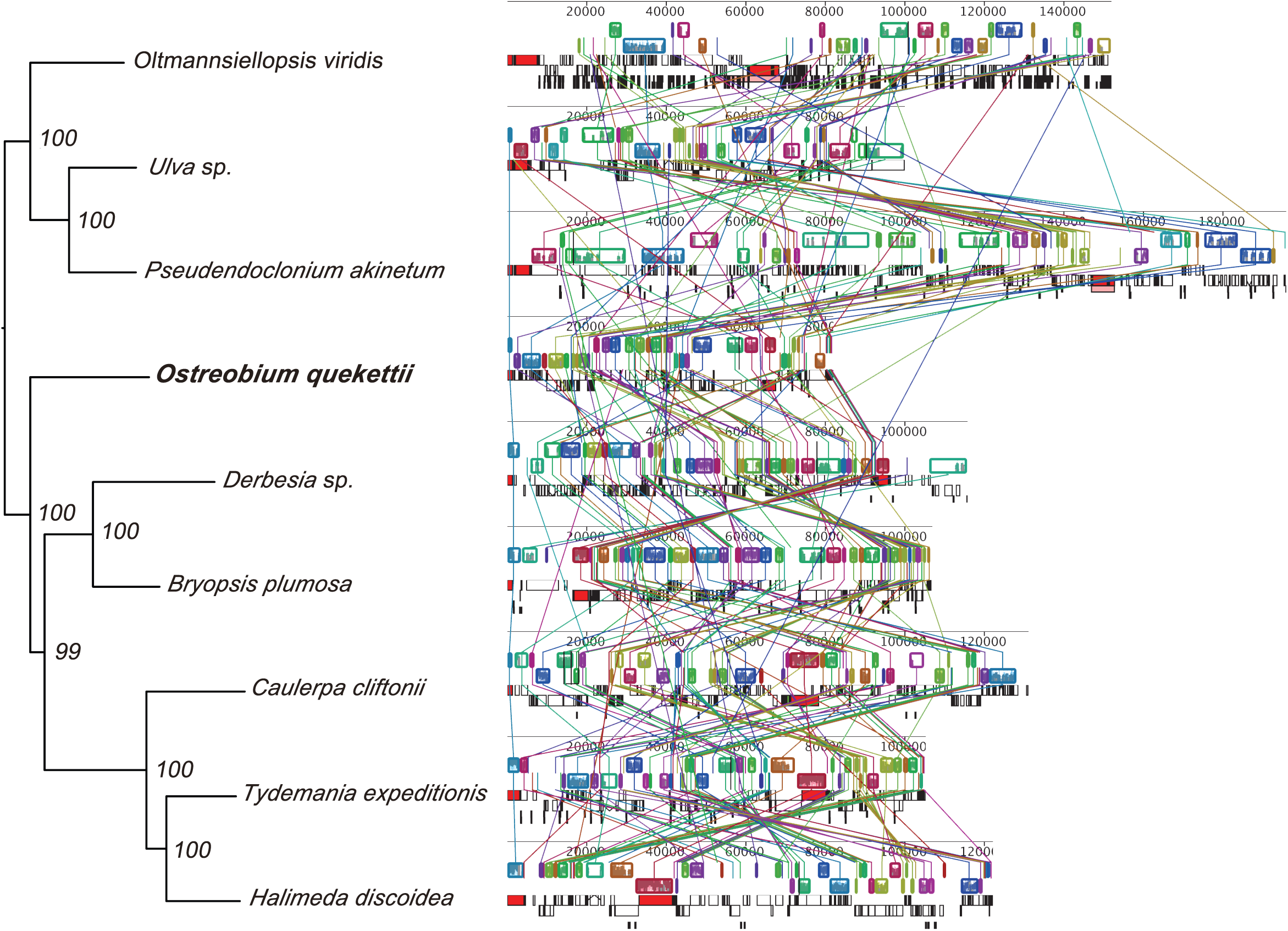
Mauve alignment of chloroplast genomes available for algae of the class Ulvophyceae, including the endolithic alga *Ostreobium quekettii* and the three seaweeds sequenced in this study. Coloured boxes indicate regions of syntheny (collinear blocks, identified by the Mauve algorithm). The species are sorted according to a Maximum Likelihood phylogeny based on a concatenated alignment of the coding sequences of the chloroplast genomes; bootstrap values are indicated near branch nodes.

### Genome economics

In order to evaluate some of our expectations regarding light-driven resource limitations on chloroplast genomes, we compared the chloroplast genome of *Ostreobium* with those of the eight other algae from the class Ulvophyceae in terms of size, compactness and nitrogen content. With 81,997 bp., *Ostreobium* has the smallest and most compact (gene-dense) chloroplast genome of all Ulvophyceae sequenced to date (Table 1, Figure 1). The size reduction in the *Ostreobium* chloroplast genom is not caused by gene loss (78 of 79 common plastid genes are present, Supplementary table 1) but by a reduction of intergenic spacers, introns and repeats (Table 1). Intergenic spacers compose only 14.9% of the *Ostreobium* plastome, compared to an average of 25.9% (std 8.2%) in other Ulvophyceae. *Ostreobium* also has the smallest number of introns, missing even the highly conserved tRNA-Leu (uaa) group I intron (Simon et al. 2003) that is present in other Bryopsidales chloroplast genomes (Leliaert and Lopez-Bautista 2015; this study). Nitrogen utilization in *Ostreobium* did not differ substantially from other algae, either in the nucleotide composition of the complete chloroplast DNA, the chloroplast transcriptome, or the amino acids of predicted proteins (Table 1). We also tested whether the average length of coding sequences is smaller in the chloroplast genome of *Ostreobium*, as gene size reduction has been observed in some endosymbionts with reduced genome sizes (e.g. Charles et al. 1999). We found that the average gene length in *Ostreobium* is similar to related algae (Supplementary table 2).

**Table 1:** Summary of the chloroplast genome features of *Ostreobium quekettii* and comparison with other Ulvophyceae plastomes.

| Species | Genome size (bp) | N content Genome | N content Transcriptome | N content Proteome | GC content (%) | Introns | Tandem repeats<sup>a</sup> | Intergenic spacers (%)<sup>b</sup> | Accession number |
| --- | --- | --- | --- | --- | --- | --- | --- | --- | --- |
| *Oltmannsiellopsis viridis* | 151,933 | 3.702 | 367.50 | 1.361 | 40.5 | 10 | 5 | 39.6 | NC_008099 |
| *Pseudendoclonium akinetum* | 195,867 | 3.6575 | 368.50 | 1.379 | 31.5 | 28 | 22 | 37.5 | AY835431 |
| *Ulva sp.* | 99,983 | 3.6265 | 367.80 | 1.366 | 25.3 | 5 | 2 | 22.7 | KP720616 |
| ***Ostreobium quekettii*** | **81,997** | **3.656** | **369.10** | **1.361** | **31.9** | **3** | **1** | **14.9** | **ERS1133536** |
| *Bryopsis plumosa* | 106,859 | 3.65 | 368.70 | 1.359 | 30.8 | 13 | 1 | 20.4 | NC_026795 |
| *Derbesia sp.* | 115,765 | 3.6445 | 366.80 | 1.375 | 29.7 | 9 | 5 | 21.4 | soon |
| *Caulerpa cliftoni*<sup>c</sup> | 131,051 | 3.688 | 365.30 | 1.376 | 37.6 | 10 | 7 | 27.0 | soon |
| *Halimeda discoidea*<sup>c</sup> | 122,140 | 3.6535 | 363.70 | 1.363 | 32.2 | 13 | 11 | 20.1 | soon |
| *Tydemania expeditionis* | 105,200 | 3.668 | 364.30 | 1.377 | 32.8 | 11 | 1 | 18.7 | NC_026796 |

<sup>a</sup> Only tandem repeats with 15bp -1000bp were included in the count.
<sup>b</sup> Excluding ORFs < 300bp.
<sup>c</sup> *Caulerpa* and *Halimeda* have, respectively, one and two scaffolds for which the number of bases is estimated but could not be determined with certainty.

### Rates of Evolution

To investigate whether the molecular pacemaker along the branch leading to *Ostreobium* is slower than in the remainder of the tree, we constructed a Maximum Likelihood (ML) phylogeny from the chloroplast genomes (71 genes concatenated, 47.559bp, Figure 1) and fitted two models of molecular evolution to the same dataset. We found that a model with differential rates of evolution for the branch leading to *Ostreobium* and the remaining branches of the phylogeny fits the data much better (ΔAIC = 92) than a model with a homogeneous rate across the entire tree. The ML parameter estimates indicate that the rate of molecular evolution along the branch leading to *Ostreobium* is 19% slower than along the other branches of the phylogeny.

### Selection on genes related to photosynthesis

Our third expectation was that genes related to the photosynthetic pathway have experienced positive selection in the lineage leading to *Ostreobium*. We concatenated genes coding different subunits of the same protein into 15 alignments based on gene classes (as defined in Wicke et al. 2011) to improve signal from short gene alignments. Two approaches were used to evaluate selection: a branch model (Yang 1998) and a random effects branch-site model (Kosakovsky Pond et al. 2011). The branch model tests whether the ω ratio (*d*_N_/*d*_S_) of a lineage specified a priori (here the branch leading to *Ostreobium*) differs from the background ω for other lineages in the phylogeny. If they do differ significantly, and if ω is greater than 1, then positive selection can be inferred for that lineage (Yang 1998). We found no indication that genes in the branch leading to *Ostreobium* have been under positive selection (Table 2). Instead, the results indicate that some proteins have a stronger signature of purifying selection in the *Ostreobium* lineage than in other branches of the phylogeny (Table 2).

**Table 2:** Omega values (*d*_N_/*d*_S_) for the different gene classes in the chloroplast genome of Ulvophycean algae. Two models were tested – a single ω for all lineages and a model with different ω values for *Ostreobium* and all other species. The goodness of fit of the two-? over the single-ω model is given by ΔAIC.

| | Single-ω model | Two-ω model | | $\Delta AIC$ |
| --- | --- | --- | --- | --- |
| | $\omega$ global | $\omega$ background | $\omega$ *Ostreobium* | |
| *Photosynthetic light reactions* |  |  |  |  |
| *atp* | 0.025 | 0.028 | 0.009 | 25.043 |
| *pet* | 0.032 | 0.034 | 0.016 | 3.625 |
| *psa* | 0.019 | 0.021 | 0.007 | 29.851 |
| *psb* | 0.029 | 0.031 | 0.013 | 40.538 |
| *Photosynthetic dark reactions* |  |  |  |  |
| *chl* | 0.022 | 0.024 | 0.012 | 4.749 |
| *ccsA* | 0.025 | 0.025 | 0.032 | -1.938 |
| *rbcL* | 0.020 | 0.023 | 0.007 | 13.922 |
| *Translation and protein-modifying enzymes* |  |  |  |  |
| *clp* | 0.014 | 0.017 | 0.004 | 2.825 |
| *infA* | 0.039 | 0.044 | 0.020 | 0.660 |
| *rpl* | 0.039 | 0.040 | 0.029 | -0.643 |
| *rps* | 0.035 | 0.036 | 0.022 | 1.179 |
| *tufA* | 0.023 | 0.025 | 0.011 | 1.516 |
| *Proteins not related to photosynthesis* |  |  |  |  |
| *accD* | 0.036 | 0.041 | 0.017 | 1.379 |
| *cys* | 0.027 | 0.027 | 6.291 | -2.000 |
| *rpo* | 0.003 | 0.003 | 0.002 | -1.749 |

A second analysis with the random effects branch-site model, which can detect selection in specific branches of the tree and sites of the alignment (Kosakovsky Pond et al. 2011), confirmed that there are no signatures of positive selection along the *Ostreobium* lineage (Supplementary table 2). As in the previous analysis, the branch-site model indicates that several gene classes have experienced stronger purifying selection in the lineage leading to *Ostreobium*: many of the genes show smaller ω values in the *Ostreobium* lineage (both mean ω and ω1 – representing purifying selection) and a higher proportion of sites under the purifying selective regime (Supplementary table 3).

## Discussion

### An economical genome

The endolithic alga *Ostreobium* has an exceptionally small and compact chloroplast genome (Table 1). Energy limitation resulting from the low light niche that this alga occupies is likely to have contributed to an evolutionary reduction of the genome size. Due to the limited light available for photosynthesis, saving energy in any aspect of its cell biology including genome replication and transcription would result in a selective advantage. Introns significantly increase the costs of transcription (Lehninger et al. 1993; Castillo-Davis et al. 2002). Likewise, repeats and intergenic spacers consume resources, so these can be under selection towards reduction in energy-poor environments (e.g. Dufresne et al. 2005; Giovannoni et al. 2005).

Genome reduction however is not always a consequence of selection favoring small genome sizes. Typically, organisms with reduced genomes are obligate parasitic or symbiotic species that have experienced relaxation of selection to maintain genes that are not essential in their niche (Mira et al. 2001; Wolf and Koonin 2013). Evolution in these cases is nearly neutral, hence the reduced genomes commonly retain pseudogenes and large amounts of non-coding DNA (Mira et al. 2001). *Ostreobium*, in contrast, has a tightly packed chloroplast genome and no sign of gene size reduction, supporting the adaptive streamlining hypothesis. It should be noted, however, that not all organisms living in low light environments have reduced genome sizes. *Acaryochloris marina* is a shade specialist and has a 8.3 Mb genome, which is large for a cyanobacteria (Swingley et al. 2008; Larsson et al. 2011). In this case, a different mechanism can be speculated on: by producing chlorophyll *d*, *Acaryochloris* may not experience the same resource constraints that *Ostreobium* does, and since it occupies a relatively uncompetitive niche this cyanobacterium could be under relaxed purifying selection which might culminate in genome expansion (see Swingley et al. 2008; Larsson et al. 2011). Wolf and Koonin (2013) proposed the existence of two phases in genome evolution: an explosive innovation phase that leads to an increase in genome complexity followed by a longer reductive phase. It is possible that the *Acaryochloris* genome size reflects its recent innovation/adaptive phase while the *Ostreobium* lineage, which has occupied an endolithic low light lifestyle for more than 500 million years (Vogel and Brett 2009; Marcelino and Verbruggen submitted), possibly has been in a reductive stage over a longer timespan. The genus *Acaryochloris*, however, is much older than 500 Ma (Sánchez-Baracaldo 2015), and in order to test this hypothesis it would be necessary to know when the genus acquired chlorophyll *d* and when it transitioned to a shaded lifestyle.

Our results are restricted to the chloroplast genome. Based on microspectrophotometry estimates, *Ostreobium*’s nuclear genome (2C ≈ 0.5 pg) is moderately small among Ulvophyceae (0.1 – 6.1 pg) (Kapraun 2007). Because no nuclear genomes of Ulvophyceae have been sequenced to date, it is not currently feasible to analyse associations between low light and nuclear genome evolution. We foresee that as more complete nuclear genomes are sequenced, equivalent analyses of the impact of resource constraints on nuclear genome evolution will follow.

Regarding gene loss, the *cemA* gene, involved in the uptake of inorganic carbon into chloroplasts (Rolland 1997), is the only gene absent from *Ostreobium* but present in all other Ulvophyceae (Supplementary table 1). Knock-out experiments in *Chlamydomonas* have shown that *cemA* is not essential for life or photosynthesis, but that its disruption drastically increases light sensitivity: mutants lacking a functional *cemA* have a lower threshold level of light perceived as excessive, so they accumulate large amounts of zeaxanthin, which is a pigment that dissipates excess light as heat (Rolland 1997). Consequently, mutants are only able to grow (photoautotrophically) under low light conditions (Rolland 1997). Although the possibility that this gene has been transferred to the nucleus in *Ostreobium* cannot be completely ruled out, *cemA* is absent from a recently sequenced *Ostreobium* transcriptome. The loss of the capacity to tolerate high light would pose a strong constraint on the transition to different habitats, which provides a plausible explanation for why *Ostreobium* lineages have diversified abundantly within the endolithic niche (Sauvage et al. 2016; Marcelino and Verbruggen submitted) but are not known to have diversified out of it (i.e., given origin to non-endolithic species). Endolithic algal species are often light saturated at low light intensities; however, some experimental studies show that they are able to photoacclimate to light levels approaching full solar irradiance (see Tribollet 2008 for a review). There are high levels of cryptic diversity within endolithic green algae and it is not known which species are able to cope with higher levels of light, raising the question of whether *cemA* has been lost in other lineages of *Ostreobium* and whether they acquired other mechanisms to tolerate high light.

We expected to observe a larger proportion of nitrogen-poor molecules in the *Ostreobium* chloroplast genome for several reasons. First, low light irradiance limits the uptake of nitrogen (MacIsaac and Dugdale 1972; Cochlan et al. 1991) and it has been empirically demonstrated that *Ostreobium* growth is limited by nitrogen and phosphorous in naturally occurring concentrations (Carreiro-Silva et al. 2012). Second, absorption of nutrients may be difficult in endolithic environments due to limited circulation and thicker diffusive boundary layers (see Larkum et al. 2003). However, our results indicate that the nitrogen content in the *Ostreobium* chloroplast genome (and predicted proteome) is similar to those of other algae in the same class (Table 1). The most obvious potential explanation for this is that many seaweeds may be affected by nitrogen limitation (Vitousek and Howarth 1991; Harrison and Hurd 2001), resulting in all of the examined genomes having similar nitrogen content. Another possibility is that *Ostreobium* may have slower generation times and metabolism than the other taxa studied, and so does not need to allocate as much nitrogen to genome replication and metabolic pathways as other algae. Nitrogen limitation may lead to size reduction of nuclear genomes in plants (Kang et al. 2015), and we do observe a size reduction of the *Ostreobium* chloroplast genome in comparison to related algae. In *Prochlorococcus*, a high-light strain that contain less nitrogen in its genome, although it is not substantially different from one of the low light strains (Dufresne et al. 2005). In this case, the nitrogen availability in the water column seems to play a more important role than a restricted nitrogen-uptake ability due to light limitation. It should also be noted that partially complete heterotrophic pathways have been observed in the genome of *Prochlorococcus*, especially in the low light strains, and so they might use other sources of energy in addition to light (García-Fernández and Diez 2004), potentially mitigating any effects of irradiance on nitrogen uptake in this organism (which would explain why low light *Prochlorococcus* strains have more nitrogen in their genomes, contrary to our expectations for *Ostreobium*). A recent review (Raven et al. 2013) suggests a theoretical association between AT/GC ratios in genomes (which could culminate in a nitrogen bias) and UV irradiation, but notes that this is not commonly observed in nature because multiple other factors influencing genome content may play a more significant role than light alone.

### Slow Rates of Evolution

We found that the *Ostreobium* lineage has slower rates of molecular evolution than closely related lineages, which could be a result of its low light niche. Sunlight, including UV radiation, induces DNA damage such as mutations and recombination (Ries et al. 2000; Raven et al. 2013, Kumar et al. 2014). While this damage often gets repaired (see Boesch et al. 2011 for mechanisms), the frequency with which remaining mutations are passed through generations dictates the molecular pacemaker (Baer et al. 2007). Following this logic, low light lineages will likely have slower rates of molecular evolution than lineages living in high light conditions, as observed in *Ostreobium* and in low light strains of *Prochlorococcus* (Dufresne et al. 2005). In *Prochlorococcus*, it is likely that the loss of DNA repair genes also contributes to an increase in mutation rates in high light strains (Dufresne et al. 2005).

Sunlight also shapes evolutionary rates through environmental energy – it sustains primary productivity and ambient temperature. Energy-rich habitats are the epicenter of evolutionary change worldwide (Davies et al. 2004; Jetz and Fine 2012; Wright and Rohde 2013). This environmental energy is positively correlated to metabolic rates in many organisms (Allen et al. 2002) and the byproducts of metabolic reactions (e.g. reactive oxygen and nitrogen species) are another major source of mutations (Gillooly et al. 2005; Boesch et al. 2011). It has been proposed that more solar radiation and higher temperatures increase metabolism and growth rates, shortening generation times and increasing mutation rates (Rohde 1992). Shorter generations lead to more mutations accumulated per unit of time, so species living in high-energy habitats tend to have faster rates of molecular evolution (Bromham 2011). One could speculate that the low energy niche that *Ostreobium* occupies results in slow metabolic rates and generation times (although they are unknown for this alga), culminating in a slow molecular pacemaker. Longer generation times have been associated with slow rates of molecular evolution in tree ferns (Zhong et al. 2014), which are also shade plants (Page 2002).

### Selection in the Ostreobium plastome

We did not find evidence for positive selection on genes related to photosynthesis in the lineage leading to *Ostreobium* (Table 2, Supplementary table 3). On the contrary, photosynthesis genes seem to be under stronger purifying selection in this lineage.

*Ostreobium* is known to have several features that facilitate low light photosynthesis. It is able to produce red-shifted chlorophylls and uses an uncommon uphill energy transfer from these chlorophylls to photosystem II (Koehne et al. 1999; Wilhelm and Jakob 2006). Many of the photosynthesis-related proteins, including the light harvesting complex superfamily, are encoded in the nucleus (Green and Parson 2003), and so innovations on these genes would not be detected in our analysis. The recently sequenced nuclear genome of the seagrass *Zostera marina* revealed an expanded number of light harvesting complex B genes (Olsen et al. 2016). Like *Ostreobium*, *Zostera* is adapted to a light depleted (aquatic) niche when compared to its land plant relatives. So far we have only analysed the plastid genome of *Ostreobium* and the major light-harvesting genes are nuclear (Green and Parson 2003, Raven et al. 2013). Thus we expect that interesting findings will result for *Ostreobium* with the analysis of transcriptome and nuclear genome data.

Another scenario that may have contributed to not observing selection is that the lineage leading to *Ostreobium* could have experienced an early burst of positive selection followed by purifying selection, and such a history may go undetected in analyses. If innovations related to low light adaptation appeared early in *Ostreobium* evolution and increased its fitness, it is expected that they would be immediately followed by purifying selection – especially if the loss of the *cemA* gene caused intolerance to high-light and confined the ancestral endolithic lineage to shaded habitats (where any mutation decreasing photosynthesis performance is likely to lead to decreased fitness). This scenario provides a plausible explanation for the stronger purifying selection in the branch leading to *Ostreobium* in comparison to other branches in the phylogeny. The available tools may not have enough power to detect faint episodes of selection, particularly if the data is saturated with synonymous substitutions or if selection occurred at deep internal branches (Kosakovsky Pond et al. 2011; Gharib and Robinson-Rechavi 2013). Simulations mimicking the evolution of algal chloroplast genomes may help to characterize those methodological limitations. Finally, the power of these analyses will certainly increase as more genomic data of high and low light-adapted lineages become available.

## Conclusion

We present the chloroplast genomes of four green algae (Bryopsidales) and investigate the genomic footprints of a low light lifestyle in the endolithic *Ostreobium quekettii*. This alga has the smallest and most gene-packed chloroplast genome among Ulvophyceae, which we suggest to be an adaptation to light-related resources constraints. The molecular pacemaker is significantly slower in the phylogenetic branch leading to *Ostreobium*, consistent with a scenario where low energy levels reduce rates of molecular evolution. Unexpectedly, we observed higher levels of purifying selection in the photosynthesis-related genes in *Ostreobium* when compared to other algae. This result could be allied to an early episodic positive selection followed by a strong purifying selection, which current methods may lack the power to detect. Sequencing additional chloroplast and nuclear genomes of different *Ostreobium* lineages and other low light adapted species will help to further clarify the genomic correlates of low light adaptations.

## Acknowledgements

This work was supported by the Australian Biological Resources Study (RFL213-08), the Australian Research Council (FT110100585, DP150100705), the Holsworth Wildlife Research Endowment and the Sapere Aude Advanced grant from the Danish Council for Independent Research for the Natural Sciences. VRM and MCMC receive a University of Melbourne scholarship. The *Caulerpa* sample was collected under the DEC Flora permit 10006072. We thank John West for providing the *Derbesia* strain and Claude Payri for facilitating field work in PNG. We are thankful to Karolina Fucikova, John Raven and the members of the Verbruggen lab for valuable insights during the execution of this study and preparation of the manuscript.

